# Orchestrated metal ion repositioning defines the dynamic catalytic strategy of the essential DNA repair nuclease APE1

**DOI:** 10.64898/2026.05.11.724140

**Authors:** Leonardo F. Serafim, Susan E. Tsutakawa, Andrew S. Arvai, Bradley R. Kossmann, Anil K. Mantha, Rachel Abbotts, David M. Wilson, Sankar Mitra, John A. Tainer, Ivaylo Ivanov

## Abstract

Human AP-endonuclease 1 (APE1) is a vital enzyme in the base excision repair pathway that protects genome stability by eliminating ubiquitous abasic DNA lesions. Despite its importance and therapeutic potential, how APE1 achieves high specificity and single-metal-ion catalytic efficiency remains unclear. Here, we present a high-resolution structure of the APE1–DNA Michaelis complex coordinated with physiological cofactor Mg^2+^. Integrating this snapshot with *ab initio* molecular dynamics and metadynamics simulations reveals a novel “moving metal ion” mechanism in which Mg^2+^ undergoes orchestrated repositioning to trigger a concerted catalytic reaction, bypassing the formation of an associative pentavalent intermediate. This distinct catalytic strategy, driven by concerted active-site reorganization, enables APE1 to efficiently process damaged DNA using only one metal ion cofactor. A previously unrecognized hydrogen-bonding network couples catalytic water activation to the metal ion movement – two events that strikingly occur on opposite sides of the active site. These findings provide a blueprint for how enzymes synchronize distal active site rearrangements with transition state formation. Our results further suggest that effective AI-targeted inhibitor design should develop capacities to predict mechanistically critical non-canonical rotamers and transient hydrogen-bonding networks. Our combined findings offer a foundation for the design of inhibitors targeting APE1 overexpression in cancer.

## INTRODUCTION

Genome integrity and replication fidelity in humans are constantly challenged by thousands of apurinic/apyrimidinic (AP or abasic) sites that are generated daily through spontaneous DNA base hydrolysis and the activity of DNA glycosylases in the base excision repair (BER) pathway(1)-5). As a cornerstone of the cellular DNA damage response, BER is integral to preventing cancer, neurodegeneration, and other disorders associated with genomic instability(2,3).

The major enzyme ensuring fast and accurate repair of these AP-site damages and BER intermediates is apurinic/apyrimidinic endonuclease 1 (APE1). APE1 efficiently and specifically cleaves the DNA backbone 5′ to the lesion, generating the 3′-hydroxyl and 5′-deoxyribose phosphate (5′-dRP) groups needed by downstream BER enzymes to enable DNA synthesis and ligation to complete repair(4,5). APE1 is overexpressed in a broad spectrum of cancers, where its elevated activity has been directly linked to resistance to DNA-damaging chemotherapeutics and ionizing radiation, making it a compelling therapeutic target. Accordingly, significant efforts have been directed toward the development of small molecule APE1 inhibitors (6-13) targeting either its endonuclease or redox signaling functions, with several compounds including APX3330(6), NO.0449-0145(14), methoxamine, and gossypol derivates, advancing into clinical evaluation (15).

DNA repair nucleases classically require two or more metal ions in their active sites to stabilize the negatively charged phosphorane-like transition state associated with phosphodiester bond cleavage and to activate water molecules for nucleophilic attack on the DNA backbone(13,16-18). A hallmark of an associative S_N_2-type phosphoryl transfer reaction, a phosphorane is a type of pentavalent phosphate intermediate with approximately equidistant apical P–O bonds (~1.7 Å in length). Divalent metal ions such as magnesium (Mg^2+^), zinc (Zn^2+^) or manganese (Mn^2+^), are consequently required to efficiently stabilize the developing negative charge on the scissile phosphate during catalysis(13,17,19-21).

Whereas most DNA repair nucleases and polymerases employ a canonical two-metal-ion mechanism, APE1 has largely been described as operating with a single Mg^2+^ ion. While no APE1–DNA structure with two native metals has been captured, kinetic and theoretical studies have raised the possibility of a second metal ion transiently binding and assisting catalysis (22,23). This has led to an ongoing debate: does APE1 utilize a second, transiently bound metal to facilitate catalysis, or has it evolved a unique strategy to achieve high efficiency with a lone cofactor? Resolving this has been challenging as crystallographic studies have largely relied on surrogate metals (e.g., Mn^2+^ or Ca^2+^) or product-state structures(24-27), which may not capture the precise positioning and role of the native Mg^2+^ ion during the catalytic transition. Consequently, understanding how APE1 achieves catalysis with its native Mg^2+^ remains a key knowledge gap, with implications for both fundamental enzymology and therapeutic strategies targeting APE1 overexpression in cancer and its role in chemotherapy resistance (15).

Here, we report the first crystal structure of the APE1 Michaelis complex bound to a native Mg^2+^ ion and DNA substrate mimic. This complex incorporates an inactivating D210N mutation and DNA with a tetrahydrofuran (THF) mimic of an AP-site. Importantly, the resulting structure effectively traps Mg^2+^ within the active site and provides an unexpected view of APE1 poised for catalysis. The new structure thereby enables a detailed comparison of Mg^2+^ ion coordination between the substrate-bound and product complexes and allowed us to dissect the specific residue interactions underlying the efficiency of APE1 catalysis.

Using this structure as a foundation for advanced Born-Oppenheimer *ab initio* molecular dynamics (BO-AIMD) and metadynamics simulations, we provide a comprehensive description of the APE1 reaction pathway. Based on first-principles quantum mechanics, our approach explicitly accounts for protein dynamics, charge transfer, and polarization effects^14,18,^(28)^-28^. Notably, this detailed description of the reactive transition, which is *a priori* unattainable using static QM/MM methods, captures critically important conformational switches that underpin efficient APE1-mediated phosphodiester cleavage of abasic DNA.

Our results uncover a surprising absence of a stabilized pentavalent intermediate. Instead, they identify a synchronous mechanism triggered by a single, mobile Mg^2+^ ion. By repositioning during the reactive trajectory, this “moving metal ion” executes multiple distinct mechanistic roles: 1) substrate activation, 2) electrostatic accommodation of the transition state, 3) hydrogen bond rearrangements that facilitate catalysis, and 4) charge stabilization of the leaving group in the latter stages of the reaction. Overall, these findings redefine our understanding of APE1 nuclease catalysis and provide a dynamic blueprint for the design of selective APE1 inhibitors.

## MATERIAL AND METHODS

### Purification and crystallization of APE1^**D210N**^

Truncated human APE1 mutant D210N (missing N-terminal 40 residues) was purified as previously described (29). The mutant crystallized with 11mer dsDNA containing a centrally located THF moiety (5′-GCTAC-THF-GATCG-3′/5′-CGATCGGTAGC-3′, Midland) in 17.5% mPEG 2K, 5% LiSO_4_, 71 mM 2-mercaptoethanol, 20 mM MgCl_2,_ and 100 mM 6.5 HEPES. Crystals were cryo-protected in 17.5% mPEG 2K, 5% LiSO4, 71 mM 2-mercaptoethanol, 20 mM MgCl_2_, 40% ethylene-glycol, and 100 mM pH 6.5 HEPES. Data was collected at the SSRL 11-1 at wavelength 0.97946 Å, at 100° K, and indexed using HKL2000. Phases were determined by molecular replacement. Refinement was performed by PHENIX(30) and COOT(31). There were two APE1/DNA complexes in the asymmetric unit (asu), with only one active site Mg^2+^ observed in the A chain. In the Ramachandran analysis, 97% were in favored positions, with no outliers. Asn212 was a rotamer outlier in both complexes. Likely due to crystallographic contacts, His289 was a rotamer outlier only in the second complex (chain D). Although most of the protein is identical between the two structures, we note that a loop between residues 146-153 in the second complex (chain D) diverged significantly from the other molecule in the asu and from the nine APE1 structures deposited in the Protein Data Bank. We base this observation on the high negative FoFc when we used the canonical loop position in the refinement model. The position of this loop was 15 Å distant from buried cysteines, and this alternative position in unlikely related to the cysteine-exposing mechanism of APE1 proposed in its role as redox effector factor (Ref-1).

### Purification of recombinant proteins for enzymatic analysis

Recombinant WT APE1 protein and APE1 structural mutants (N68A, D70A, E96A) were purified as described previously (32-37). To normalize protein concentration, 250 ng of each recombinant protein was resolved by SDS-PAGE, and the resultant gels were stained using SYPRO Red [Life Technologies, Frederick MD; 1:5000 in 7.5% (v/v) acetic acid]. Protein bands were imaged and quantified using a Typhoon fluorescence imager and ImageQuant TL software (GE Healthcare, Piscataway NJ). Recombinant proteins were adjusted to final concentrations in dilution buffer [50mM HEPES-KOH (pH 7.5), 50 mM KCl, 5 mM DTT, 0.01% Triton X-100, 5% glycerol, 100 μg/ml bovine serum albumin.

### Oligonucleotide substrate

An 18-base oligonucleotide containing THF (5′-GTCACCTGT-THF-TACGACTC-3′; TriLink Biotechnologies, San Diego, CA) was radiolabeled with [γ-^32^ATP] at the 5′-end using T4 polynucleotide kinase (New England Biolabs, Ipswich MA). End-labeled oligonucleotides were annealed to the complementary unlabeled oligonucleotide in a thermocycler by heating at 95°C for 3 min and then cooling by 2°C every 2 min to 25°C.

### AP site incision assay

Recombinant protein (50 pg/150 pM or 500 pg/1.5 nM) was incubated for the indicated time in a 10 µl final volume with 1 pmol (100 nM) radiolabeled double-stranded, THF-containing 18mer oligonucleotide substrate at 37°C in 50 mM HEPES-KOH (pH 7.5), 50 mM KCl, 10 mM MgCl_2_, 0.01% Triton X-100, 100 μg/ml bovine serum albumin for 15 min. Reactions were inhibited by the addition of stop buffer (95% formamide, 20 mM ethylenediaminetetraacetic acid, 0.5% bromphenol blue, 0.5% xylene cyanol) and heated to 95°C for 5 min. Reaction products were resolved by 20% polyacrylamide urea denaturing gel electrophoresis, imaged, and quantified using a Typhoon phosphorimager and ImageQuant TL software (GE Healthcare).

### Classical Molecular Dynamics Simulations

We used our crystal structure of APE1 D210N in complex with DNA as the starting structure for the simulations of the reactant state (S_1_). Our previously published structure (PDB identification code: 4IEM)(29) was used for the simulations of the product state (S_3_). The mutation of the catalytic residue N210D was performed with UCSF Chimera (38). The THF molecule of the nucleotide inside the active site was modified by adding a hydroxyl group to model the hemiacetal form of the AP site.

The classical MD simulations were performed with the PMEMD module of the Amber software package (39). The AMBER ff14SB force field was employed for the protein, while the OL15 forcefield was employed for the DNA(40). The parameters for the nonstandard nucleotide were developed using Antechamber(39). The charges and electrostatic potentials (ESP) were computed at the HF/6-31G(d,p) level of theory, after geometry optimization at the B3LYP/6-31G(d,p) level of theory using Gaussian software (41). The enzyme-substrate complex was then placed in a cubic box of 100 x 100 x 100 Å dimensions. The minimal distance from the edge of the box to the surface of the protein complex was 15 Å. The TIP3P water model(42) was used for the solvent, and Na^+^ and Cl^-^ ions were added to neutralize the total charge of the system and mimic physiological concentration (0.154 M).

After minimization, the system was heated up to 300 K over 10 ns at constant volume (NVT) while imposing positional restraints of 100 kcal mol^-1^ Å^-2^ on the heavy atoms. Subsequently, the restraints were slowly removed, and a production MD run of 200 ns was performed in the isothermal-isobaric ensemble (NPT). The temperature control (300 K) was performed by a Langevin thermostat with a collision frequency of 1 ps^-1^, and the pressure control (1 atm) was accomplished by a Monte Carlo barostat. The SHAKE algorithm was used to constrain the bonds involving hydrogen atoms, and the particle mesh Ewald method(43) was used to compute the electrostatic interactions. A 10 Å short-range non-bonded interaction cutoff distance and a 1 fs time step were used for the simulations.

### QM/MM and Ab initio Molecular Dynamics (AIMD)

We performed QM/MM Born-Oppenheimer MD simulations using the CP2K code, which combines the QM program QUICKSTEP and the MM driver FIST (28). The electrostatic coupling between the QM and MM regions was computed using a real-space multigrid technique(44). The QM region of the system was simulated in a 20 x 20 x 20 Å box and a wavelet Poisson solver employed to decouple QM images(45). The QM subsystem comprises: the side chains of residues Asn 68, Asp 70, Glu 96, Tyr 171, Asp 210, Asn 212 and His 309, the Mg^2+^ ion, three water molecules, and the nucleotide (the phosphate and the two adjacent ribose rings). The QM region was treated at the DFT/BLYP(46) level using a dual basis set of Gaussians and plane-waves (GPW)(47). DFT-D3 dispersion correction was applied (48). We employed a double zeta (MOLOPT) basis set along with an auxiliary PW basis set with a density cutoff of 600 Ry and Goedecker-Teter-Hutter (GTH) pseudopotentials (49). Protein residues outside the QM region, DNA, ions, and solvent atoms comprised the MM subsystem and were simulated as classical particles using the AMBER ff14SB/OL15 force field(40). All AIMD simulations were performed in the NVT ensemble (300 K), employing a Nosé thermostat. A time step of 0.5 fs was employed during 100-ps production simulations. All structures were visualized with UCSF Chimera (38).

### Reaction Pathway Optimization and Metadynamics

The optimization of the reaction pathway was carried out using constrained dynamics path optimization method, within a QM/MM framework, as implemented in the CP2K software (28). The DFT functional, basis set, pseudopotentials and other QM/MM settings were consistent with those used in the AIMD simulations. The computational setup involved the use of 10 equally spaced replicas along the reaction path to represent the transition between the reactant (S_1_) and product (S_3_) states. The replicas were optimized using 3 collective variables: the nucleophile-electrophile distance (O_W_-P), the scissile nucleotide bond length (P-O_LG_) and the metal-leaving group distance (Mg-O_LG_).

The free energy profiles associated with the phosphodiester bond cleavage were obtained using metadynamics (MTD) simulations(50). The optimized structures obtained from the string method(51) were used as the starting point for the MTD simulations. In these simulations multidimensional Gaussian hills potentials with a height of 0.5 kcal/mol and a width of 0.2 Å were deposited along the respective CVs, at every 100 fs to build-up the MTD biasing potential and enhance sampling of the reactive transitions. We employed MDT in multiple-walker replica runs with 10 walkers to improve convergence of the free energy profiles along the phosphoryl transfer reaction. We used the same CVs as defined in the string method to explore the reaction pathway. Three sets of MTD simulations were performed, each using one, two, and three CVs, corresponding to one, two, and three-dimensional reaction coordinates, respectively. This approach allowed us to systematically investigate the influence of additional CVs on the free energy landscape. In aggregate, the MTD free energy simulations required 100 ps of dynamics to achieve convergence.

## RESULTS

### Crystal structure of an APE1–DNA Michaelis complex with native metal coordination

To obtain a structure of the human APE1–DNA–Mg substrate complex, we crystallized a catalytically inactive D210N APE1 mutant with a centrally located AP site mimetic tetrahydrofuran (THF) and Mg^2+^ (**Fig. 1** and **Supplementary Table 1**). The structure is significant for providing the first view of the APE1 Michaelis complex with native metal ion coordinated in the active site. Asp210 is critical for APE1’s catalytic activity and has been posited as the general base that performs water activation prior to the nucleophilic attack on the scissile phosphate. Asp210 is located on the opposite side of the active site from the Mg^2+^ position in the product complex. Thus, a D210N mutation does not directly interfere with Mg^2+^ binding, allowing an unprecedented view of the changes in magnesium coordination between the substrate and product complexes. The structure was determined at a 2.05 Å resolution with an R and R-free values of 0.18 and 0.23, respectively. We did not observe electron density consistent with an attacking water molecule in the active site, suggesting that a D210N substitution inactivates the enzyme by displacing the catalytic water molecule.

**Figure 1.**
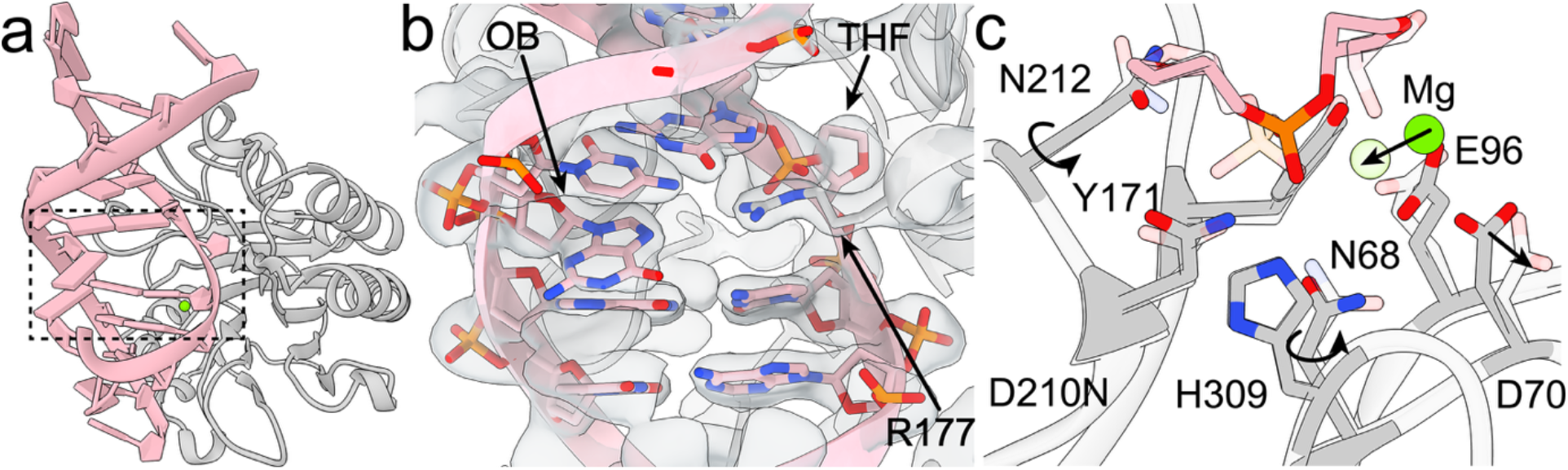
High-resolution crystal structure of APE1 D210N Michaelis complex with native metal coordination. (a) Overall structure of the enzyme-substrate complex with APE1 (gray) and 11-mer DNA (pink). (b) Close-up view of the THF nucleotide inserted into the active site and the orphan base (OB) paired with Arg177 with electron density (gray surface). (c) Detailed view of the active site in the APE1 D210N enzyme-substrate complex (solid color) overlaid with the product structure (PDB ID: 4IEM, shown with transparency). Key side-chain translations and rotations, and metal displacement are indicated with arrows.

The Mg^2+^ binding site in the APE1 Michaelis complex features water-mediated coordination to the scissile phosphate of the abasic site and differs from all previous APE1 structures (**Fig. 1**). Notably, active site overlays of the D210N mutant and product structures (PDB ID: 4IEM)(29) show distinct metal coordination geometries. In the product complex, Mg^2+^ is directly coordinated to the DNA backbone via the phosphate groups flanking the cleavage site. The metal is also coordinated by residue Glu96. The increased number of direct DNA-metal contacts in the product implies tighter metal coordination upon completion of the APE1 reaction. By contrast, in the Michaelis complex the Mg^2+^ ion is shifted by 2.3 Å relative the product and coordinates DNA indirectly via a structural water molecule (**Fig. 1c**). Predictably, metal coordination is much weaker in the substrate complex. Correspondingly, Mg^2+^ was observed in only one of the two chains in the asymmetric unit of the crystal.

Notably, we observe unexpected conformational shifts of the active site residues in the D210N mutant structure (**Fig. 1c**) relative to the product (PDB ID: 4IEM) and apoenzyme (PDB ID: 4QHD)(52) structures. These conformational changes have key implications for catalysis and involve: 1) a ~120° Asn68 side chain rotation, 2) a shift of the Asp70 residue by ~1.4 Å towards Asn68, 3) a ~60° side chain rotation of the metal coordinating Glu96 residue, and 4) a ~120° side chain rotation of the catalytically essential Asn212 residue. Side chain rotations were confirmed by hydrogen-bond distances with neighboring oxygen/nitrogen atoms and by stronger density at the oxygen position of the asparagine side chains. The Mg^2+^ ion did not appear to alter side chain orientations, as the RMSD was 0.082 Å between residues Asp70, Glu96, Tyr171, Asp210, Asn212 and His309 in chain A and chain D, whose coordinates were refined without NCS.

The observed active site changes between reactant and product are driven by a previously unrecognized hydrogen bond network, extending on both sides of the scissile phosphate from the general base (Asp210) toward the magnesium binding site. The network encompasses residues Asn212, Asp210, Asn68, Asp70 and Glu96 (**Supplementary Fig. 1**). Importantly, the D210N mutant is a structural mimic of the catalytic intermediate that would form after activation of the attacking nucleophilic water molecule. Water activation leads to a protonated Asp210. Removal of the negative charge at this position (as in the D210N mutant) appears to disrupt hydrogen bonding between Asn68 and Asp210 and may cause the Asn68 side chain to flip and interact with Glu96. In turn, movement of Asp70 triggers a concomitant shift of the metal-coordinating residue Glu96. In the wild-type (WT) product structure, Asn68 is coordinated to water molecules from the first coordination shell of Mg^2+^, preventing Asn68 rotation. Our findings further suggest that the observed changes in the hydrogen bonding pattern may impact the switch from weak to strong Mg^2+^–DNA association between reactant and product. Notably, the newly identified hydrogen bonding network provides a suitable relay mechanism to alter the Mg^2+^ coordination geometry in response to the evolving charge on the Asp210 general base during the water activation step of the catalytic mechanism. In sum, perturbing the hydrogen-bonding network across the active site in the D210N mutant reveals a network of interactions with an apparent role in dynamically choreographing the catalytic steps in the APE1 mechanism.

Comparison to the previously reported APE1 structure with a phosphothiolate DNA substrate and non-native Mn^2+^ ion (PDB ID: 5DG0) is also instructive (**Supplementary Fig. 2**). In the 5DG0 structure the metal ion occupies a position shifted by ~1.8 Å relative to Mg^2+^ in our structure. Superposition of the active sites also reveals pronounced conformational differences: residue Asn212 undergoes a ~40° rotation, Asn68 rotates by ~170°, Glu96 by ~30°, and Asp70 shifts by ~1.4 Å. These changes collectively reshape the coordination environment and modulate interactions between the metal ion, catalytic residues, and the scissile phosphate. Similar rearrangements are observed in the phosphothiolate substrate structure lacking a metal ion (PDB ID: 5DFI) and in the E96Q/D210N double mutant (PDB ID: 5DFJ), both showing ~40° rotation of Asn212 and ~170° rotation of Asn68.

### Kinetic analysis confirms Asn68 importance for the incision reaction

If proton abstraction during water activation by Asp210 leads to Asn68 rotation and a concurrent shift in magnesium coordination, it follows that mutating Asn68 should significantly impact APE1 catalysis. Indeed, steady-state reactions with a human APE1 N68A mutant revealed an ~100-fold reduction of incision activity relative to the WT protein (**Fig. 2** and **Supplementary Table 2**). Notably, this is a 5- to 10-fold stronger effect compared to the catalytic impairment caused by directly mutating the two metal coordinating residues of the apoenzyme and product structures (Glu96 and Asp70). Specifically, E96A mutation results in ~20-fold reduction of catalytic activity whereas D70A mutation incurs an even more modest ~10-fold reduction. N68A did not exhibit as strong of an effect on activity as mutations of residues Asp210 (10,000-fold decrease) or Asn212 (7000-fold decrease), which directly position and activate the nucleophilic water molecule in the APE1 mechanism. In terms of its deleterious effect on catalysis, N68A substitution falls in-between these two cases. The reduction of specific activity upon N68A mutation is consistent with the Asp68 rotation being important for the APE1 catalytic reaction.

**Figure 2.**
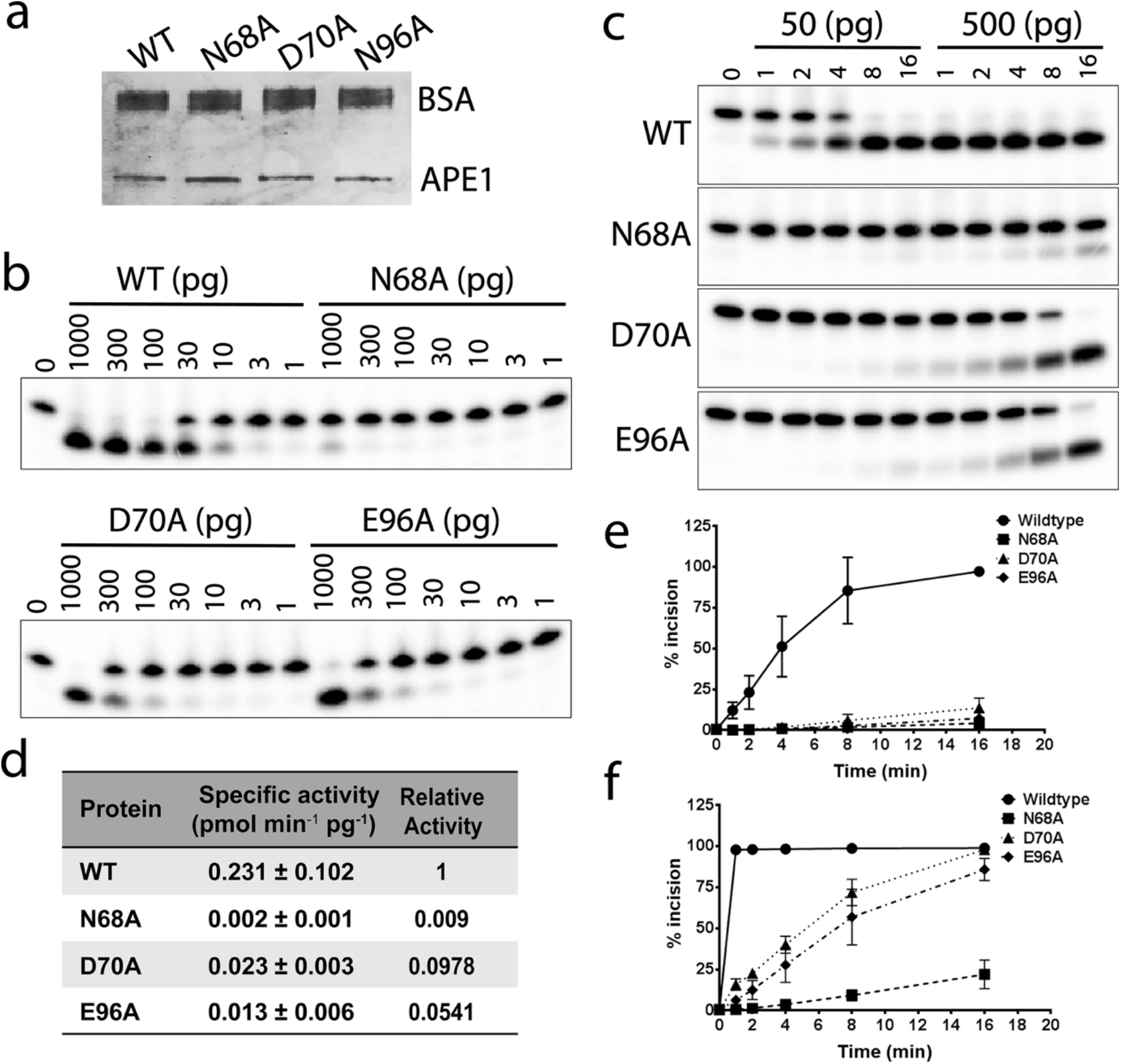
Critical biochemical roles for Asn68, Asp70, and Glu96 in APE1 catalysis. (a) APE1 protein purity and comparative concentration. Purified APE1 proteins (250 ng, as indicated) were separated on an SDS-PAGE and stained with SYPRO red. BSA was added to the sample for comparative purposes. (b) APE1 concentration-dependent incision activity. Indicated amounts of APE1 protein (WT or designated site-specific mutant) were incubated with P32-labeled 18-mer F-DNA duplex substrate (1 pmol) for 5 min in 10 μl volumes, followed by electrophoresis on a standard 20%-urea PAG to resolve the initial substrate (top band) from the incised product (bottom band). (c) Representative gels showing time-dependent incision activity of APE1 proteins. Incision reactions were performed at two APE1 amounts (50 and 500 pg) for the designated times (0, 1, 2, 4, 8, and 16 min). See panel b for further details. (d) Specific activities of APE1 proteins. Values were generated from >3 independent experimental data points and are presented relative to WT APE1 activity. (e and f) Incision activity plots for APE1 proteins. Data represents incision efficiency at 50 pg protein (e) or 500 pg protein (f) at the indicated time point.

### Classical molecular dynamics of APE1 D210N supports a moving Mg2+ ion mechanism

To test whether the change in the Mg^2+^ coordination sphere was attributable only to the D210N mutation or induced by the dissimilar binding of an intact AP-DNA substrate versus 3′-hydroxyl and 5′-deoxyribose DNA fragments, we performed classical molecular dynamics (MD) simulations of the APE1 (D210N) crystal structure with bound Mg^2+^ and substrate DNA. In the models, a protonated aspartic acid was introduced in place of the mutated N210 residue. Additionally, a hydroxyl group was incorporated in the THF structure to recreate the natural AP-site structure.

We tested three metal positions: one from the highest resolution product structure (PDB ID: 4IEM), one from the APE1 (D210N) structure, and one from a DNA-free structure (PDB ID: 4LND). In all three structures, the Mg^2+^ ions are in the same area of the active site and are separated by 1-2 Å from one another.

In the product structure, the Mg^2+^ ion is coordinated only to Glu96, while in the apoenzyme, WT and D210N structures, Mg^2+^ is coordinated to both Glu96 and Asp70. The MD simulation of the WT model with the side chains rotated as in the APE1 (D210N) mutant resulted in unstable Mg^2+^ coordination, while the models of the product and the apoenzyme with side chain placements retained the Mg^2+^ in its original position.

These results are in line with the hypothesis that the rotations and shifts observed in the APE1 (D210N) crystal structure would interfere with binding of Mg^2+^ at the apoenzyme Mg^2+^ position resulting in its movement during catalysis. Moreover, in the MD simulations of the APE1 (D210N) mutant and WT APE1 with protonated D210 the nucleophilic water molecule was not retained in place, rationalizing the absence of electron density in our crystal structure. This instability is associated with the hydrogen bonding network formation between Asn212, Asp210 and Asn68. Notably, the change in hydrogen bonding pattern induced by negative charge removal at residue D210 restricts the available space to accommodate a water in position suitable for catalysis (**Supplementary Fig. 3**). Thus, we posit that the APE1 mechanism requires a metal that can move during catalysis and that this motion is coupled to the side chain flipping of the Asn68 residue.

### BO-AIMD modeling reveals catalytically essential active-site conformational switching

To provide a comprehensive description of the APE1 catalytic mechanism, we first created QM/MM models(44) of the ternary APE1–Mg^2+^–DNA complexes in the substrate (S_1_) and product (S_3_) states and simulated them using *ab initio* molecular dynamics(28,44). In the substrate complex (S_1_), Asp210 serves as a general base for catalysis and is suitably positioned within a short hydrogen-bonding distance (*d*_*HB*_ = 2.75 ± 0.08 Å)^*^ from the nucleophilic water molecule (**Fig. 3a, Supplementary Movie 1**). In this position, Asp210 is oriented to activate the water molecule and promote a nucleophilic attack on the phosphorus atom of the AP-site. Correspondingly, the average interatomic distance between the activated water oxygen and the phosphorus atom of the scissile phosphate (O_w_–P bond) is 3.75 ± 0.20 Å. Additionally, the angle between the electrophilic phosphorus atom and the apical oxygens of the nucleophile and the leaving group (O_w_–P–O_LG_ angle) was 153.2 ± 5.8°. Both values are optimal for initiating the DNA cleavage reaction. Asn212 is another key catalytic residue proximal to Asp210 that helps orient the nucleophilic water molecule. Mutation of Asn212 practically abrogates APE1 catalysis. Yet, at this early stage of the mechanism Asn212 orientation is still suboptimal (positioned >3.7 Å away from the water molecule). We further show that the Asn212 side chain reorients in the intermediate state S_2_ to assist catalysis.

**Figure 3.**
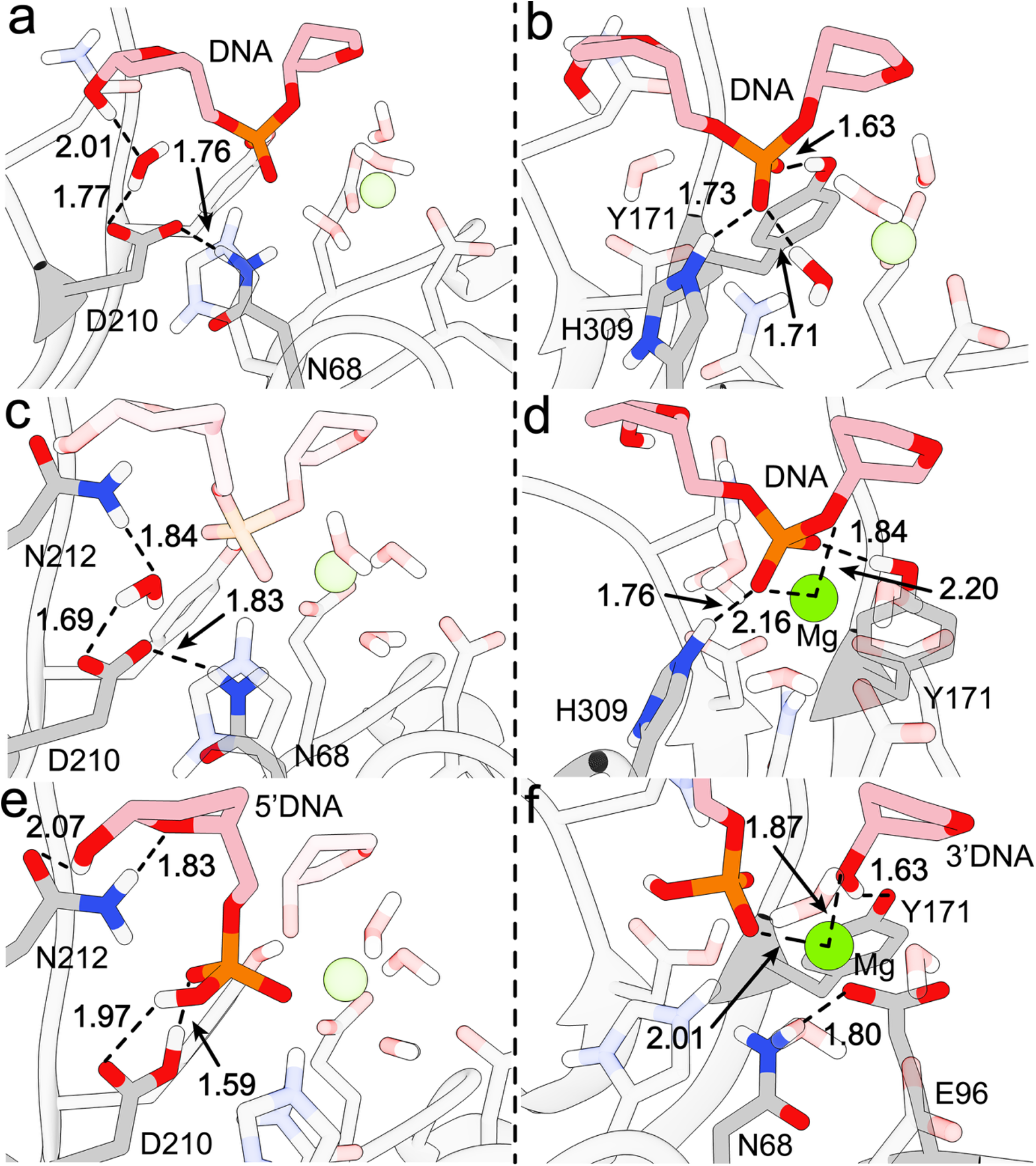
Concerted conformational shifts of Asp210, Asn212, Asn68, and Glu96 revealed by BO-AIMD are crucial for the APE1 excision mechanism. (a, b) Michaelis complex (state S1), highlighting interactions at the water activation site (a) and the metal binding site (b). (c, d) Intermediate state S2, showing (c) side-chain rotation of Asn212 and (d) coordination of Mg^2+^ to two oxygen atoms of the scissile phosphate group. (e, f) Product state S3, showing hydrogen bond interactions stabilizing the 5′-phosphate fragment (e) and Mg^2+^ interactions anchoring the 3′-hydroxyl fragment within the active site (f).

Two catalytically essential residues, Tyr171 and His309, were found to form stable hydrogen bonds with the equatorial oxygen atoms of the scissile phosphate at average distances of 2.67 ± 0.11 Å and 2.75 ± 0.14 Å, respectively. During MD equilibration, residue His309 was modelled as protonated based on our pKa calculations (computed pKa = 8.78) and an earlier NMR study (53). The BO-AIMD simulations confirmed this assignment as His309 retained its positive charge despite being able to proton transfer in a QM/MM setting. Tyr171 and His309 not only anchor the scissile phosphate in the Michaelis complex but are also crucial for stabilizing the developing negative charge on the phosphate group during catalysis (**Fig. 3b**). The Mg^2+^ ion flanking the scissile phosphate, is coordinated to Asp70, Glu96 and three water molecules. Notably, in state S_1_ there is no significant Lewis acid activation of the DNA substrate by Mg^2+^, as the closest phosphate oxygen atom is >4.0 Å away from the metal ion. In state S_1_, the key catalytic residue Asn68 forms a strong hydrogen bond (*d*_*HB*_ = 2.79 ± 0.14 Å) with the putative general base residue Asp210.

BO-AIMD simulations of the product state (S_3_) reveal a rearranged APE1 active site, accompanied by significant changes in the hydrogen bonding pattern. The rearrangement features a 2.5-Å shift in the Mg^2+^ ion position and a switch from water-mediated to direct coordination of the metal. Specifically, in S_3_ the magnesium ion binds to both the non-bridging oxygen of the AP-site phosphate and the oxygen atom of the leaving group (**Fig. 3d, Supplementary Movie 1**). Movement of the metal ion is accompanied by the dissociation of the coordinating Asp70 residue and a concerted ~120° rotation of the Asn212 side chain. Additionally, in S_3_ the 5′-dRP fragment is held by hydrogen bonds from the phosphate to Asp210 (*d*_*HB1*_ = 3.16 ± 0.27 Å and *d*_*HB2*_ = 2.67 ± 0.15 Å) while Asn212 forms hydrogen bonds with the AP-site ribose ring (*d*_*HB1*_ = 3.00 ± 0.27 Å and *d*_*HB2*_ = 3.01 ± 0.11 Å). Concurrently, protonation and charge neutralization of Asp210 induces flipping of the Asn68 side chain, switching its hydrogen bonding pattern. In turn, the switch in Asn68-mediated hydrogen bonding couples the water activation event to conformational changes in the metal binding site. Additionally, the phosphate anchoring residue Tyr171 shifts from coordinating the non-bridging oxygen atom of the phosphate group to hydrogen bonding with the oxygen atom of leaving group (*d*_*HB*_ = 2.63 ± 0.15 Å). The observed reorientation suggests a dual role for Tyr171: 1) stabilizing the phosphorane-like transition state during the nucleophilic attack; and 2) neutralizing of the leaving group at the end of the reaction.

The intermediate state S_2_ was obtained by path optimization connecting states S_1_ and S_3_. The path optimization allowed us to resolve subtle rearrangements accompanying Mg^2+^ reorientation. In S_2_, the direct Mg^2+^ ion contact provides the necessary Lewis acid activation to stabilize the developing negative charge on the scissile phosphate group in the transition state of the phosphoryl transfer reaction (**Fig. 3e, Supplementary Movie 1**). The scissile P–O_LG_ bond lengthens from 1.62 Å in S_1_ to 1.86 Å in S_2_, confirming the metal-induced polarization of the leaving-group bond. Importantly, Asn212 undergoes a conformational shift to provide essential hydrogen bonding to the oxygen atom of the AP-site ribose ring (*d*_*HB*_ = 3.19 ± 0.29 Å) and the lesion specific hydroxyl group (*d*_*HB*_ = 3.02 ± 0.22 Å), while establishing a strong interaction with the nucleophilic water molecule (*d*_*HB*_ = 2.94 ± 0.21 Å). Collectively, these observations indicate that S_2_ represents the near attack conformation in which the active-site geometry is organized to trigger hydrolysis.

### A moving Mg^2+^ ion fulfils disparate mechanistic roles in early-versus late-stage catalysis

The overarching complexity of the APE1 reaction necessitates the use of a composite reaction coordinate. This involves not only the atoms directly participating in phosphoryl transfer (O_w_, P and O_LG_) but also concomitant metal coordination shifts, Asn212 and Asn68 side chain rotations, hydrogen bond rearrangements and proton transfer events that accompany catalysis. This task cannot be accomplished using intuitive reaction coordinates or static QM/MM models. Instead, we relied on chain-of-replicas path optimization in the space of multiple collective variables. Specifically, we combined constrained *ab initio* molecular dynamics (AIMD) with a variant of the string method (SM)(51) to compute a minimum free energy path (MFEP) for the APE1 phosphoryl transfer reaction. The MFEP path represents the entire catalytic mechanism, including all on-path intermediates and transition states. Notably, the MFEP also identifies the dominant motions of the active site residues that ensure transition state stabilization in response to changing active site environment during catalysis.

Using this methodology, we connected the S_1_ and S_3_ states and identified a distinct intermediate state (S_2_) that is formed prior to hydrolysis of the phosphate bond. State S_2_ represents a key stage of the reaction involving activation of substrate DNA (**Fig. 3c**). Formation of intermediate S_2_ involves Asn212 side chain rotation and a Mg^2+^ ion shift from indirect to direct coordination at the AP-site (**Supplementary Movie 2**). The activation free energy DG^≠^ of this mechanistic step (associated with transition state T_1_) is 11.0 kcal/mol. Therefore, this is not the rate limiting step of the overall reaction (**Fig. 4** and **Fig. 5**). However, the Lewis acid effect, and the polarization of the scissile phosphate bond provided by divalent metal ion binding in S_2_ are essential for the subsequent hydrolytic cleavage of DNA. Thus, contrary to previous mechanistic proposals for APE1(54), the DNA cleavage reaction catalyzed by APE1 requires direct Mg^2+^ ion coordination to the scissile phosphate, which sets up the correct geometry for inline water-mediated nucleophilic attack.

**Figure 4.**
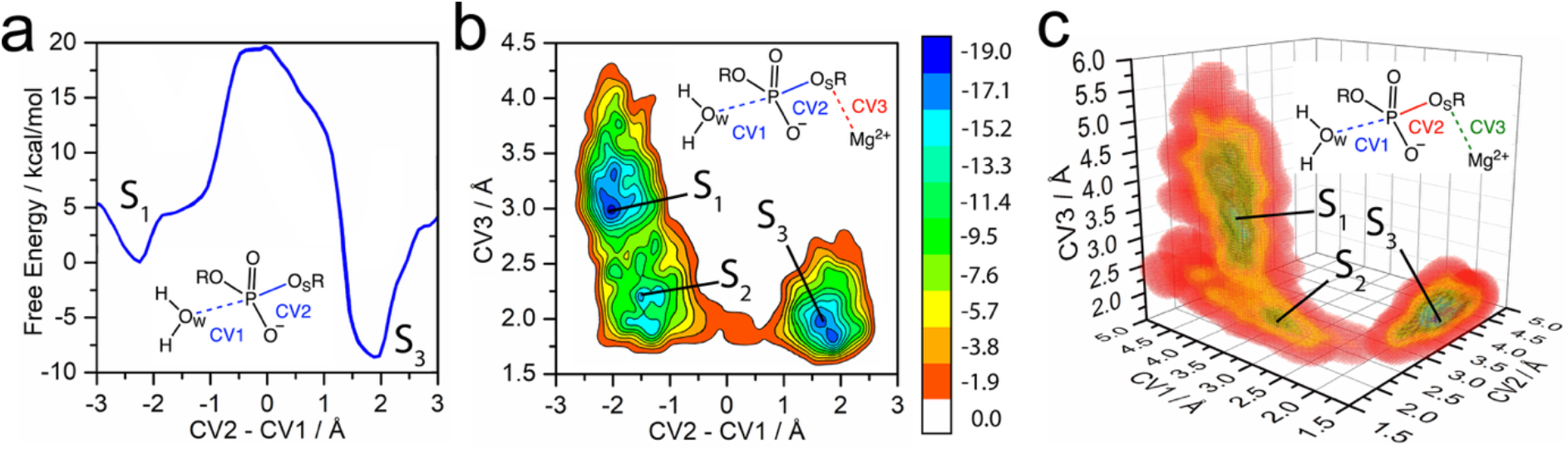
The optimized reaction pathway from ab initio QM/MM supports a concerted S_N_2-type mechanism without a stabilized phosphorane intermediate. Free energy surfaces obtained by metadynamics simulations using one (a), two (b), and three (c) collective variables. Graphical representations of the collective variables used to build the MTD biasing potential are shown as insets. S_1_, S_2_ and S_3_, represent the Michaelis-Menten complex, intermediate and product states of APE1 catalysis (as depicted in **Fig. 3**).

**Figure 5.**
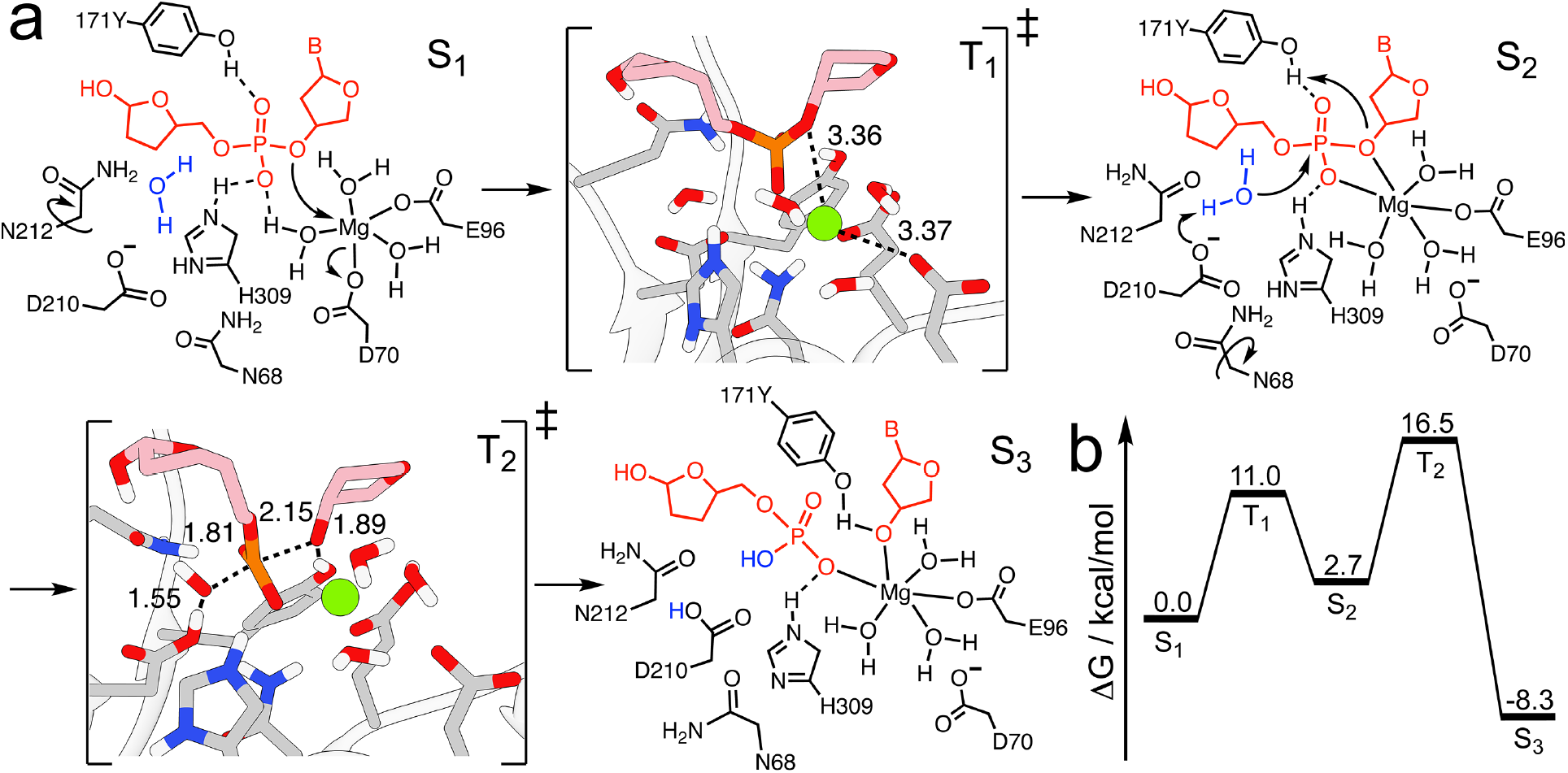
Unified APE1 catalytic mechanism schematic. (a) Representative structures of each of the stable states (S_1_, S_2_ and S_3_) and the transition states (T_1_ and T_2_). The representative structures of T_1_ and T_2_ were extracted from the MTD trajectories based on the minima of the free energy surfaces obtained by *ab initio* metadynamics with 3 collective variables (**Fig. 4c**). (b) Free energy diagram showing energy barriers associated with the S_1_ → S_2_ → S_3_ transition pathway.

Transition state T_2_ connects states S_2_ and S_3_. During the reaction step, Asp210 abstracts a proton from the nucleophilic water molecule for an attack on the phosphorus atom (**Supplementary Movie 3**). This is followed by the formation of a S_N_2-like transition state and a pyramidal inversion at the phosphorus atom, eventually leading to dissociation of the leaving group oxygen atom (O_LG_). The 2.15 Å distances between the apical oxygen atoms and the electrophilic phosphorus in state T_2_ are not compatible with the expected bond distances in a stabilized phosphorane intermediate (~1.7 Å), thus precluding a fully associative mechanism. Instead, we posit that the reactive transition occurs via a concerted A_N_D_N_ type mechanism (55).

Besides the metal ion, residues Tyr171 and His309 are critical for stabilization of the negative charge on the scissile phosphate in transition state T_2_. Specifically, His309 engages a phosphate equatorial oxygen atom via a stable hydrogen bond and contributes a full positive charge toward transition state stabilization: this mimics the effect of a coordinating second metal ion, as also consistent with comparative superpositions with the three-metal-ion bacterial abasic site endonuclease IV^29^. On the opposite side of the scissile phosphate, Tyr171 initially hydrogen bonds to an equatorial oxygen atom, providing electrostatic stabilization of T_2_ through charge transfer and polarization effects. Thus, the transition state of the rate limiting step in the APE1 mechanism is triply stabilized by positive charges: 1) the divalent magnesium ion, 2) the positively charged His309, and 3) the charge on Tyr171 OH. The triple stabilization effect results in an activation free energy DG^≠^ of 16.5 kcal/mol for this mechanistic step (**Fig. 5**) that is tied to the overall rate of the nuclease reaction and explains the exceptional catalytic proficiency of APE1. The computed TS barrier is in close agreement with experimentally measured catalytic rate for WT APE1(56). By contrast, a computational attempt to transition directly from S_1_ to S_3_ without the active site rearrangements identified through path optimization, result in an artificially elevated reaction barrier of ~35 kcal/mol.

After the pyramidal inversion, Tyr171 switches coordination from the equatorial oxygen atom to the apical O_LG_ oxygen atom of the leaving group. The presence of a strong hydrogen bond and the closely matched pK_a_ values of the tyrosine and the leaving group result in proton sharing and partial neutralization of the negative charge on O_LG_, which is also stabilized by divalent metal binding. Indeed, in the BO-AIMD simulations we observe frequent proton hopping events between the Tyr171 oxygen atom and O_LG_, suggestive of a low-barrier, proton-sharing hydrogen bond between these functional groups. Thus, we posit that Tyr171 plays two distinct, yet critical roles in the APE1 mechanism: 1) initial stabilization of the equatorial oxygen, and 2) subsequent partial charge transfer and proton-sharing with the leaving group.

## DISCUSSION

The canonical view of APE1 catalysis presupposes a static active site, pre-formed upon the steric recognition and “sculpting” of the abasic site into a shallow, damage-specific pocket. Yet, the detailed catalytic mechanism of APE1 has remained largely elusive and controversial due to the absence of high-resolution structures capturing both the Michaelis complex and the product state with intact amino acid configurations and native metal ions. Consequently, existing mechanistic models fail to explain how APE1 achieves its remarkable catalytic efficiency with a single metal ion or how it prevents off-target genomic incisions that would be lethal to the cell. Our results provide the missing structural information to redefine the APE1 catalytic paradigm. Integrated structural and computational data show that APE1 catalysis does not rely on a static architecture. Instead, it utilizes an atypical synchronized, “moving single-metal-ion” mechanism (**Fig. 5**). This non-canonical strategy enables APE1 to perform complex chemistry using a single metal ion cofactor, differing from the multi-metal-ion mechanisms common to most DNA repair nucleases(13,16-18).

By explicitly accounting for active site dynamics, our findings uncover critical conformational rearrangements and functional roles for all first- and second-shell residues experimentally implicated as key for catalysis. Importantly, we thereby resolve controversies regarding the general base identity and the role of Tyr171. While a Y171F mutation was associated with a ~1,200-fold reduction in catalytic efficiency, our computational analyses refute the hypothesis that Tyr171 can serve as a general base or directly engage the nucleophilic water through hydrogen bonding (54). Here BO-AIMD simulations indicate that Tyr171 is spatially incapable of hydrogen-bonding to the nucleophile without cleaving the DNA on the incorrect side of the AP-site. Instead, we identify Asp210 as the general base, working in tandem with Asn212 to orient and deprotonate the water molecule for an in-line nucleophilic attack. Within this mechanistic framework, Tyr171 is reassigned two noncanonical roles: 1) stabilization of the developing negative charge in the transition state and 2) protonation of the 3′-oxygen of the leaving group. Importantly, this “division of labor” between Asp210 and Tyr171 explains how the enzyme achieves precise phosphodiester cleavage while maintaining substrate specificity.

BO-AIMD QM/MM simulations further elucidate the coordinated choreography of active site dynamics and moving metal ion interactions during the APE1 catalytic cycle (**Supplementary Movie 2 and Movie 3**). A striking feature of the newly defined APE1 reaction is thus the dynamic repositioning of the Mg^2+^ ion. We find that the Mg^2+^ ion evolves through three distinct functional states (**Fig. 5**).

In the reactant state (S_1_) the Mg^2+^ ion establishes monodentate coordination to Asp70 and Glu96 and coordinates three water molecules. During the transition from state S_1_ to S_2_ the metal ion repositions to engage an equatorial oxygen atom of the scissile phosphate. This event triggers Asp210 and Asn212 side chain rotations, and Asp70 dissociation from Mg^2+^. Such conformational rearrangements emphasize the dynamic nature of the single metal-ion-mediated APE1 catalysis. Interestingly, formation of the first transition state (T_1_) involves the transient neutralization of Asp210 and His309, mediated by the nucleophilic water molecule, resulting from a rapid His309 side chain shift in response to metal binding, as both groups bind the same oxygen atom. This orchestrated active-site reorientation prepares APE1 for nucleophilic attack by stabilizing the water nucleophile and generating a reactive configuration.

In S_2_, the Mg^2+^ ion directly binds the scissile phosphate, providing Lewis acid activation and lowering the free energy barrier of the hydrolysis reaction by ~15 kcal/mol (compared to an S1-initiated reaction). The S_2_ to S_3_ transition begins with Asp210 abstracting a proton from the nucleophilic water molecule. This enables the resulting hydroxide ion to attack the electrophilic phosphorus atom in a concerted S_N_2 reaction step. Our analysis does not support the formation of a stabilized phosphorane intermediate. This observation is consistent with a single-metal-ion mechanism, in which a single Mg^2+^ ion can electrostatically support the developing negative charge in the transition state without the need to stabilize a pentavalent intermediate. In contrast, enzymes that employ two or three metal ions often stabilize such intermediates as part of their catalytic pathways. The additional metals in these systems provide enhanced electrostatic stabilization and additional coordination sites to support the intermediate’s trigonal bipyramidal geometry.

In S_3_ Tyr171 forms a proton donor hydrogen bond with the same oxygen atom of the leaving group that is now also associated with the Mg^2+^ ion. The role of the Mg^2+^ ion evolves from structural stabilization of the Michaelis complex (S_1_) to Lewis acid activation of the electrophilic phosphorus atom in state S_2_ and eventually to charge neutralization of the leaving group in state S_3_. The repositioning of the metal ion plus the Asn68, Asp210 and Asn212 side chain rotations underscore the dynamic nature of APE1 catalysis. Tyr171 plays a dual role in stabilizing the substrate in the transition from S_1_ to S_2_ and contributes to leaving group neutralization in the S_2_ to S_3_ transition, rather than a metal ion bound water, as posited in previous reports(54). These findings also challenge previous hypotheses by showing that Asp210 and Asn212 rather than Asp210 and Tyr171 are responsible for positioning and activation of the nucleophile. Moreover, the absence of a phosphorane intermediate highlights the efficiency of single metal ion catalysis in APE1, distinguishing it from the canonical multimetal systems including AP-endonuclease endonuclease IV (54)(57).

APE1’s repair activity in tumor cells antagonizes both radiotherapy and chemotherapy. Thus, novel APE1 inhibitors are urgently needed for clinical applications. Rational design and modification of inhibitor molecules critically depend on integrated understanding of structure and catalytic mechanism, as exemplified by our study. Given that APE1 overexpression is a hallmark of poor prognosis in multiple cancers, our findings provide a structural and mechanistic blueprint for the design of selective, mechanism-based inhibitors.

Moreover, for inhibitor design, the extraordinary malleability of the APE1 active site presents a significant challenge to current AI-based protein structure predictions. APE1 is an active target for preclinical and clinical cancer drug discovery efforts (9-13) including AI approaches(58). Yet, current AI models such as AlphaFold3, which prioritize static structures and thermodynamic minima, fail to predict the non-canonical rotamers and transient hydrogen-bonding networks identified here (**Supplementary Fig. 2d**). Thus, our results have implications for AI-Driven discovery and inhibitor design, suggesting APE1 as a benchmark for such developments.

In our APE1 analysis, we uncover the specificity and efficiency of moving metal ion catalysis, identify the “hotspots” of metal repositioning, and define the choreography for active site residue dynamic evolution through three distinct functional states. Together these findings provide a dynamic blueprint for the design of selective inhibitors. Such therapeutics would provide a promising approach to counteract APE1 overexpression in treatment-resistant cancers, turning a fundamental mechanistic insight for DNA repair biology into viable strategies for clinical intervention.

## Supporting information

Combined Supplementary Figures and Tables

## AUTHOR CONTRIBUTIONS

I.I. and J.A.T. directed the study. L.F.S., S.E.T., J.A.T. and I.I. contributed to the design of the study. L.F.S. performed the model building and molecular simulations. L.F.S., A.A., T.P., R.A., S.E.T., J.A.T., D.W., A.K.M. and I.I. performed experiments and/or analyzed the data. L.F.S., T.P., R.A., S.E.T., J.A.T., D.W., A.K.M., S.M., and I.I. wrote the manuscript.

## SUPPLEMENTARY DATA STATEMENT

Supplementary Data are available at NAR online.

## DATA AVAILABILITY

Structural models of the APE1 complexes in the S_1_, S_2_ and S_3_ states have been deposited in the ModelArchive database with corresponding accession codes: 1) ma-1k1tm [https://modelarchive.org/doi/10.5452/ma-1k1tm]; 2) ma-po7jq [https://modelarchive.org/doi/10.5452/ma-po7jq]; 3) ma-ogeri [https://modelarchive.org/doi/10.5452/ma-ogeri]. The structure of the human APE1–DNA– Mg^2+^ ternary complex carrying a D210N mutation was deposited in the PDB with accession code: 9Z4Y [https://www.rcsb.org/structure/9Z4Y]. PDB accession codes of all structures referenced in this study: 4IEM [https://www.rcsb.org/structure/4IEM], 4QHD [https://www.rcsb.org/structure/4QHD], 5DG0 [https://www.rcsb.org/structure/5DG0], 5DFI [https://www.rcsb.org/structure/5DFI], 5DFJ [https://www.rcsb.org/structure/5DFJ], 4LND [https://www.rcsb.org/structure/4LND].

## Footnotes

* average bond lengths and angles are presented with corresponding standard deviation (s. d.) values throughout.

## REFERENCES

1. Lindahl, T. (1993) Instability and decay of the primary structure of DNA. Nature, 362, 709–715.

2. Tubbs, A. and Nussenzweig, A. (2017) Endogenous DNA damage as a source of genomic Instability in cancer. Cell, 168, 644–656.

3. Ayyildiz, D., Antoniali, G., D’Ambrosio, C., Mangiapane, G., Dalla, E., Scaloni, A., Tell, G. and Piazza, S. (2020) Architecture of the human APE1 interactome defines novel cancers signatures. Sci Rep, 10, 28.

4. Whitaker, A.M. and Freudenthal, B.D. (2018) APE1: A skilled nucleic acid surgeon. DNA Repair, 71, 93–100.

5. Liu, Y., Prasad, R., Beard, W.A., Kedar, P.S., Hou, E.W., Shock, D.D. and Wilson, S.H. (2007) Coordination of steps in single-nucleotide base excision repair mediated by apurinic/apyrimidinic endonuclease 1 and DNA polymerase beta. J Biol Chem, 282, 13532–13541.

6. Caston, R.A., Gampala, S., Armstrong, L., Messmann, R.A., Fishel, M.L. and Kelley, M.R. (2021) The multifunctional APE1 DNA repair-redox signaling protein as a drug target in human disease. Drug Discov Today, 26, 218–228.

7. Xue, Z. and Demple, B. (2022) Knockout and inhibition of APE1: Roles of APE1 in base excision DNA repair and modulation of gene expression. Antioxidants (Basel), 11,1817.

8. Laev, S.S., Salakhutdinov, N.F. and Lavrik, O.I. (2017) Inhibitors of nuclease and redox activity of apurinic/apyrimidinic endonuclease 1/redox effector factor 1 (APE1/Ref-1). Bioorg Med Chem, 25, 2531–2544.

9. Wilson, D.M., 3rd, Deacon, A.M., Duncton, M.A.J., Pellicena, P., Georgiadis, M.M., Yeh, A.P., Arvai, A.S., Moiani, D., Tainer, J.A. and Das, D. (2021) Fragment- and structure-based drug discovery for developing therapeutic agents targeting the DNA Damage Response. Prog Biophys Mol Biol, 163, 130–142.

10. Hu, M., Zhang, Y., Zhang, P., Liu, K., Zhang, M., Li, L., Yu, Z., Zhang, X., Zhang, W. and Xu, Y. (2025) Targeting APE1: Advancements in the diagnosis and treatment of tumors. Protein Pept Lett, 32, 18–33.

11. Sharma, A., Grimsley, H.E., Courtemanche, K. and Powell, S.N. (2025) Contrasting roles of APE1 and APE2 in genome maintenance, cancer development, and therapeutic targeting. NAR Cancer, 7, zcaf048.

12. Torrado, C., Gonzalez-Ortiz, A., Xavier, C.B., Tan, H.N., Ngoi, N.Y.L. and Yap, T.A. (2025) Drugging the DNA damage response in the clinic: going beyond PARP. Expert Rev Anticancer Ther, 1–16.

13. Ivanov, I., Tainer, J.A. and McCammon, J.A. (2007) Unraveling the three-metal-ion catalytic mechanism of the DNA repair enzyme endonuclease IV. Proc Nat Acad Sci U S A, 104, 1465–1470.

14. Long, K., Gu, L., Li, L., Zhang, Z., Li, E., Zhang, Y., He, L., Pan, F., Guo, Z. and Hu, Z. (2021) Small-molecule inhibition of APE1 induces apoptosis, pyroptosis, and necroptosis in non-small cell lung cancer. Cell Death & Disease, 12, 503.

15. Malfatti, M.C., Bellina, A., Antoniali, G. and Tell, G. (2023) Revisiting two decades of research focused on targeting APE1 for cancer therapy: The pros and cons. Cells, 12, 1895

16. Yang, W. (2011) Nucleases: diversity of structure, function and mechanism. Q Rev Biophys, 44, 1–93.

17. Steitz, T.A. and Steitz, J.A. (1993) A general two-metal-ion mechanism for catalytic RNA. Proc Nat Acad Sci U S A, 90, 6498–6502.

18. Lassila, J.K., Zalatan, J.G. and Herschlag, D. (2011) Biological phosphoryl-transfer reactions: understanding mechanism and catalysis. Annu Rev Biochem, 80, 669–702.

19. Serafim, L.F., Jayasinghe-Arachchige, V.M., Wang, L., Rathee, P., Yang, J., Moorkkannur N S. and Prabhakar, R. (2023) Distinct chemical factors in hydrolytic reactions catalyzed by metalloenzymes and metal complexes. Chem Commun, 59, 8911–8928.

20. Yang, W., Lee, J.Y. and Nowotny, M. (2006) Making and breaking nucleic acids: two-Mg2+-ion catalysis and substrate specificity. Mol Cell, 22, 5–13.

21. Ivanov, I. and Klein, M.L. (2005) Dynamical flexibility and proton transfer in the arginase active site probed by ab initio molecular dynamics. J Am Chem Soc, 127, 4010–4020.

22. Beernink, P.T., Segelke, B.W., Hadi, M.Z., Erzberger, J.P., Wilson, D.M. and Rupp, B. (2001) Two divalent metal ions in the active site of a new crystal form of human apurinic/apyrimidinic endonuclease, APE1: implications for the catalytic mechanism. Edited by I. A. Wilson. J Mol Biol, 307, 1023–1034.

23. Oezguen, N., Schein, C.H., Peddi, S.R., Power, T.D., Izumi, T. and Braun, W. (2007) A “moving metal mechanism” for substrate cleavage by the DNA repair endonuclease APE-1. Proteins, 68, 313–323.

24. Freudenthal, B.D., Beard, W.A., Cuneo, M.J., Dyrkheeva, N.S. and Wilson, S.H. (2015) Capturing snapshots of APE1 processing DNA damage. Nat Struct Mol Biol, 22, 924–931.

25. Weaver, T.M., Hoitsma, N.M., Spencer, J.J., Gakhar, L., Schnicker, N.J. and Freudenthal, B.D. (2022) Structural basis for APE1 processing DNA damage in the nucleosome. Nat Commun, 13, 5390.

26. Whitaker, A.M., Flynn, T.S. and Freudenthal, B.D. (2018) Molecular snapshots of APE1 proofreading mismatches and removing DNA damage. Nat Commun, 9, 399.

27. Mol, C.D., Izumi, T., Mitra, S. and Tainer, J.A. (2000) DNA-bound structures and mutants reveal abasic DNA binding by APE1 DNA repair and coordination. Nature, 403, 451–456.

28. Kühne, T.D., Iannuzzi, M., Del Ben, M., Rybkin, V.V., Seewald, P., Stein, F., Laino, T., Khaliullin, R.Z., Schütt, O., Schiffmann, F. et al. (2020) CP2K: An electronic structure and molecular dynamics software package - Quickstep: Efficient and accurate electronic structure calculations. J Chem Phys, 152, 194103.

29. Tsutakawa, S.E., Shin, D.S., Mol, C.D., Izumi, T., Arvai, A.S., Mantha, A.K., Szczesny, B., Ivanov, I.N., Hosfield, D.J., Maiti, B. et al. (2013) Conserved structural chemistry for incision activity in structurally non-homologous apurinic/apyrimidinic endonuclease APE1 and endonuclease IV DNA repair enzymes. J Biol Chem, 288, 8445–8455.

30. Afonine, P.V., Grosse-Kunstleve, R.W., Echols, N., Headd, J.J., Moriarty, N.W., Mustyakimov, M., Terwilliger, T.C., Urzhumtsev, A., Zwart, P.H. and Adams, P.D. (2012) Towards automated crystallographic structure refinement with Phenix.Refine. Acta Crystallogr D, 68, 352–367.

31. Casañal, A., Lohkamp, B. and Emsley, P. (2020) Current developments in Coot for macromolecular model building of Electron Cryo-microscopy and Crystallographic Data. Protein Sci, 29, 1069–1078.

32. Erzberger, J.P., Barsky, D., Schärer, O.D., Colvin, M.E. and Wilson, D.M., III. (1998) Elements in abasic site recognition by the major human and Escherichia coli apurinic/apyrimidinic endonucleases. Nucleic Acids Res, 26, 2771–2778.

33. Erzberger, J.P. and Wilson, D.M., 3rd. (1999) The role of Mg2+ and specific amino acid residues in the catalytic reaction of the major human abasic endonuclease: new insights from EDTA-resistant incision of acyclic abasic site analogs and site-directed mutagenesis. J Mol Biol, 290, 447–457.

34. Nguyen, L.H., Barsky, D., Erzberger, J.P. and Wilson, D.M., 3rd. (2000) Mapping the protein-DNA interface and the metal-binding site of the major human apurinic/apyrimidinic endonuclease. J Mol Biol, 298, 447–459.

35. Hadi, M.Z., Ginalski, K., Nguyen, L.H. and Wilson, D.M., 3rd. (2002) Determinants in nuclease specificity of Ape1 and Ape2, human homologues of Escherichia coli exonuclease III. J Mol Biol, 316, 853–866.

36. Hadi, M.Z., Coleman, M.A., Fidelis, K., Mohrenweiser, H.W. and Wilson, D.M., 3rd. (2000) Functional characterization of Ape1 variants identified in the human population. Nucleic Acids Res, 28, 3871–3879.

37. Kim, W.C., Berquist, B.R., Chohan, M., Uy, C., Wilson, D.M., 3rd and Lee, C.H. (2011) Characterization of the endoribonuclease active site of human apurinic/apyrimidinic endonuclease 1. J Mol Biol, 411, 960–971.

38. Pettersen, E.F., Goddard, T.D., Huang, C.C., Couch, G.S., Greenblatt, D.M., Meng, E.C. and Ferrin, T.E. (2004) UCSF Chimera--a visualization system for exploratory research and analysis. J Comput Chem, 25, 1605–1612.

39. Salomon-Ferrer, R., Case, D.A. and Walker, R.C. (2013) An overview of the Amber biomolecular simulation package. WIREs Comput Mol Sci, 3, 198–210.

40. Cheatham III, T.E. and Case, D.A. (2013) Twenty-five years of nucleic acid simulations. Biopolymers, 99, 969–977.

41. Frisch, M.J., Trucks, G.W., Schlegel, H.B., Scuseria, G.E., Robb, M.A., Cheeseman, J.R., Scalmani, G., Barone, V., Petersson, G.A., Nakatsuji, H. et al. (2016) Gaussian 16 Rev. C.01., Wallingford, CT.

42. Jorgensen, W. L., Chandrasekhar, J., Madura, J. D., Impey, R. W., & Klein, M. L. (1983). Comparison of simple potential functions for simulating liquid water. J Chem Phys, 79, 926–935.

43. Darden, T., York, D. and Pedersen, L. (1993) Particle mesh Ewald: An N·log(N) method for Ewald sums in large systems. J Chem Phys, 98, 10089–10092.

44. Laino, T., Mohamed, F., Laio, A. and Parrinello, M. (2005) An efficient real space multigrid QM/MM electrostatic coupling. J Chem Theory Comput, 1, 1176–1184.

45. Blöchl, P.E. (1995) Electrostatic decoupling of periodic images of plane-wave-expanded densities and derived atomic point charges. J Chem Phys, 103, 7422–7428.

46. Lee, C., Yang, W. and Parr, R.G. (1988) Development of the Colle-Salvetti correlation-energy formula into a functional of the electron density. Phys Rev B, 37, 785–789.

47. VandeVondele, J., Krack, M., Mohamed, F., Parrinello, M., Chassaing, T. and Hutter, J. (2005) Quickstep: Fast and accurate density functional calculations using a mixed Gaussian and plane waves approach. Comput Phys Commun, 167, 103–128.

48. Grimme, S., Antony, J., Ehrlich, S. and Krieg, H. (2010) A consistent and accurate ab initio parametrization of density functional dispersion correction (DFT-D) for the 94 elements H-Pu. J Chem Phys, 132, 154104.

49. VandeVondele, J. and Hutter, J. (2007) Gaussian basis sets for accurate calculations on molecular systems in gas and condensed phases. J Chem Phys, 127, 114105.

50. Barducci, A., Bonomi, M. and Parrinello, M. (2011) Metadynamics. WIREs Computational Molecular Science, 1, 826–843.

51. e, W., Ren, W. and Vanden-Eijnden, E. (2002) String method for the study of rare events. Physical Review B, 66, 052301.

52. He, H., Chen, Q. and Georgiadis, M.M. (2014) High-resolution crystal structures reveal plasticity in the metal binding site of apurinic/apyrimidinic endonuclease I. Biochemistry, 53, 6520–6529.

53. Lowry, D.F., Hoyt, D.W., Khazi, F.A., Bagu, J., Lindsey, A.G. and Wilson, D.M. (2003) Investigation of the role of the histidine–aspartate pair in the human exonuclease III-like abasic endonuclease, APE1. J Mol Biol, 329, 311–322.

54. Aboelnga, M.M. and Wetmore, S.D. (2019) Unveiling a single-metal-mediated phosphodiester bond cleavage mechanism for nucleic acids: A multiscale computational investigation of a human DNA repair enzyme. J Am Chem Soc, 141, 8646–8656.

55. Casalino, L., Nierzwicki, Ł., Jinek, M. and Palermo, G. (2020) Catalytic mechanism of non-target DNA cleavage in CRISPR-Cas9 revealed by ab initio molecular dynamics. ACS Catalysis, 10, 13596–13605.

56. DeHart, K.M., Hoitsma, N.M., Thompson, S.H., Borin, V.A., Agarwal, P.K. and Freudenthal, B.D. (2025) APE1 active site residue Asn174 stabilizes the AP-site and is essential for catalysis. J Biol Chem, 301, 110655.

57. Garcin, E.D., Hosfield, D.J., Desai, S.A., Haas, B.J., Bjoras, M., Cunningham, R.P. and Tainer, J.A. (2008) DNA apurinic-apyrimidinic site binding and excision by endonuclease IV. Nat Struct Mol Biol, 15, 515–522.

58. Iqbal, A.B., Masoodi, T.A., Bhat, A.A., Macha, M.A., Assad, A. and Shah, S.Z.A. (2025) Explainable AI-driven prediction of APE1 inhibitors: enhancing cancer therapy with machine learning models and feature importance analysis. Mol Divers, 29, 3371–3390.

