## Supplementary material for "Orchestrated metal ion repositioning defines the dynamic catalytic strategy of the essential DNA repair nuclease APE1": Combined Supplementary Figures and Tables

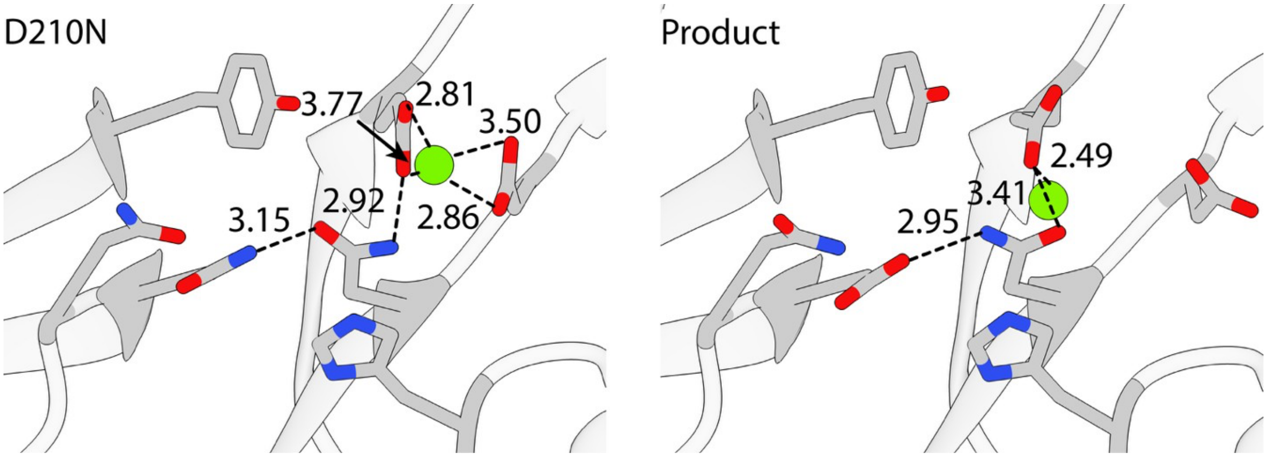


**Supplementary Figure 1. Comparison of key active-site contacts in the APE1–DNA Michaelis complex and product states.** Key residue interactions in the D210N APE1–DNA Michaelis complex structure (left) and the WT APE1–DNA product structure (right, PDB ID: 4IEM). These structures are overlaid in **Figure 1c** of the main manuscript. Interatomic distances are given in Å. Distinct hydrogen-bonding patterns between the reactant and product states highlight the reorganization of catalytic residues and DNA backbone geometry during the phosphodiester cleavage reaction.

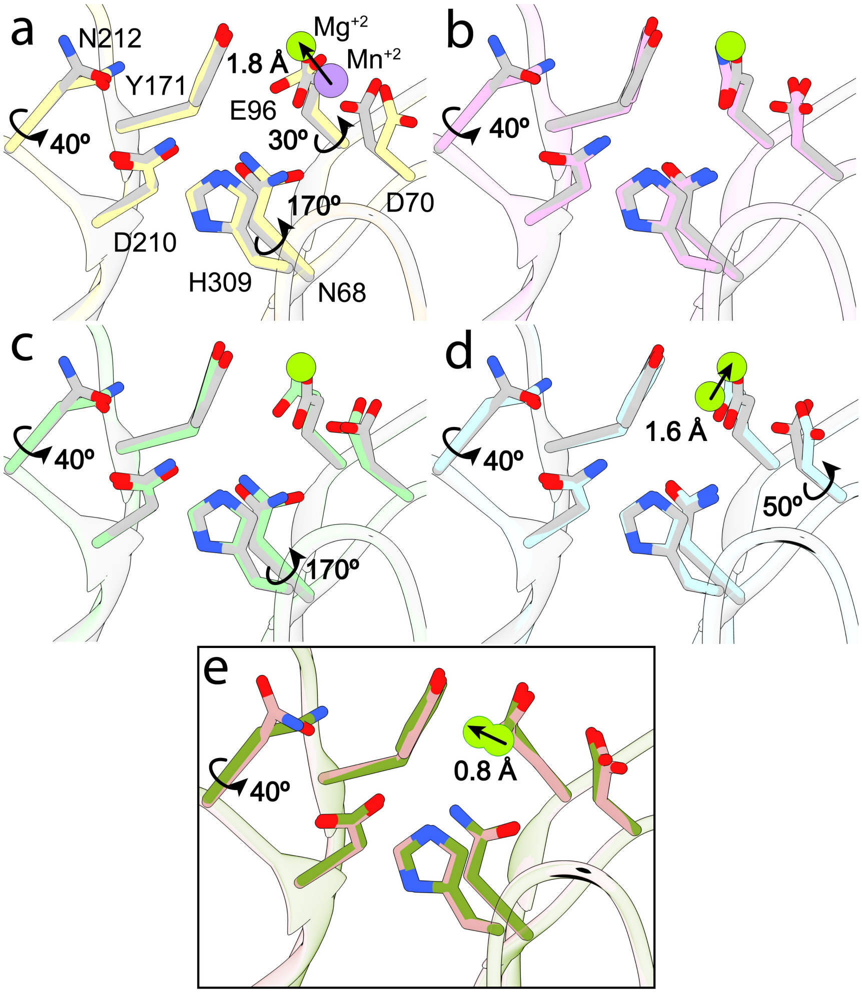


**Supplementary Figure 2. Structural comparison of the D210N APE1–DNA–$Mg^{2+}$ structure with previously reported phosphorothiolate substrate structures.** Overlays of the D210N APE1–DNA–$Mg^{2+}$ structure with: (a) the APE1-phosphorothiolate complex containing a non-native $Mn^{2+}$ ion (PDB ID: 5DG0); (b) the phosphorothiolate complex lacking a bound metal ion (PDB ID: 5DFI); and (c) the E96Q/D210N double-mutant structure without metal (PDB ID: 5DFJ). Relative to the $Mg^{2+}$-bound Michaelis complex, these structures exhibit substantial rearrangements in the metal-binding environment, including shifts in metal position (in 5DG0) and large rotations of key catalytic residues (Asn212 and Asn68) that alter coordination geometry and interactions with the scissile phosphate. In (d), our new structure of the Michaelis complex is overlayed with a corresponding AlphaFold3 model. In (e), AlphaFold3 model for the product state is overlayed with our previously reported product-state structure (4IEM). Notably, AlphaFold3 does not reproduce the characteristic residue movements associated with the metal-bound catalytic state.

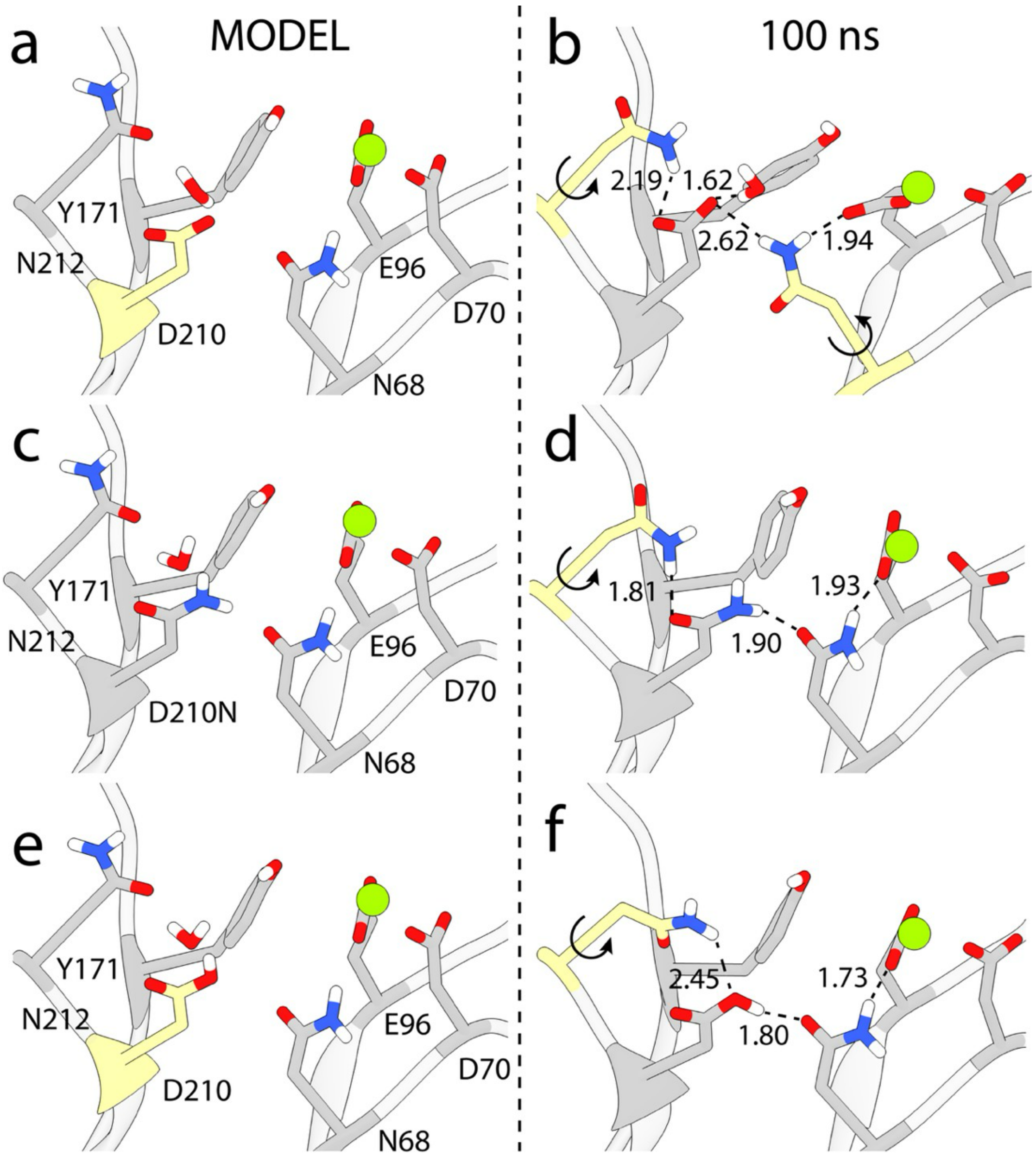


**Supplementary Figure 3. Initial models and equilibrated classical MD structures derived from the D210N APE1–DNA structure.** Panels (a), (c), and (e) show the starting structures, and panels (b), (d), and (f) show the corresponding configurations after 100 ns of classical MD simulations. (a) Wild-type model generated from the D210N APE1–DNA structure with Asp210 in its deprotonated (negatively charged) state; (c) the D210N APE1–DNA mutant structure; (e) wild-type model generated from the same structure but with Asp210 protonated. Panels (d) and (f) illustrate that a neutral residue at position 210 cannot maintain or accommodate a catalytic water molecule within the active site.

#### SUPPLEMENTARY TABLES

| | APE1 D210N-$Mg^{2+}$-DNA |
|---|---|
| PDB Code | 9Z4Y |
| Wavelength | 0.97946 |
| Resolution range | 35.9 - 2.05 (2.12-2.05) |
| Space group | P 1 21 1 |
| Unit cell a, b, c (Å) a, {3, y (°) | 47.0, 128.5, 64.8<br>90, 96.4, 90 |
| Total reflections | 170760 |
| Unique reflections | 46722 (4116) |
| Multiplicity | 3.7 (2.6) |
| Completeness (%) | 93.0 (53.54) |
| Mean I/sigma(I) | 9.9 (2.3) |
| Wilson B-factor | 28.2 |
| R-meas | 11.3 (75.2) |
| CC1/2 | 87.8(94.1) |
| Reflections used in refinement | 44179 (1453) |
| Reflections used for R-free | 2212 (67) |
| R-work | 0.18 |
| R-free | 0.23 |
| Number of non-hydrogen atoms | 5696 |
| -Protein | 4376 |
| -DNA | 854 |
| -Ligand/ion | 25 |
| -Water | 441 |
| RMS (bonds) | 0.003 |
| RMS (angles) | 0.69 |
| Ramachandran favored (%) | 97.1 |
| Ramachandran allowed (%) | 2.9 |
| Ramachandran outliers (%) | 0.0 |
| Rotamer outliers (%) | 0.4 |
| Clashscore | 0.5 |
| Average B-factor (all) | 33 |
| -macromolecules only | 30 |
| -ligands only | 43 |
| -solvent only | 36 |

**Supplementary Table 1. X-ray data collection and refinement statistics for the APE1 D210N-$Mg^{2+}$-DNA complex.** Crystals of truncated human APE1 D210N bound to an 11-mer dsDNA containing a tetrahydrofuran (THF) abasic-site analog were obtained in 17.5% mPEG 2K, 5% $LiSO_4$, 71 mM 2-mercaptoethanol, 20 mM $MgCl_2$, and 100 mM HEPES pH 6.5. Data were collected at SSRL beamline 11-1 (/\ = 0.97946 Å) at 100 K and processed with HKL2000. Phases were determined by molecular replacement, and structures were refined using PHENIX and COOT. The asymmetric unit contains two APE1/DNA complexes, with a single active-site $Mg^{2+}$ observed in chain A. Refinement statistics, geometry validation, and model composition are summarized below.

| Protein | Binding | Incision |
|---|---|---|
| Wild-Type | 1 | 1 |
| N68A | 1§ | 0.009# |
| D70A | 1 | 0.1 |
| D70R | 1 | 0.04 |
| E96A | 1 | 0.05# |
| E96Q | 1 | 0.0005 |
| Y128A | <0.01 | 0.25 |
| R156Q | <0.01 | 0.01 |
| Y171F | 1§ | 0.0002 |
| Y171H | 1§ | 0.00006 |
| D210A | 1 | <0.00004* |
| D210N | 1§ | <0.00004* |
| D210H | 1§ | 0.0006 |
| F266A | >0.15 | 0.18 |
| D283N | n.d. | 0.1§ |
| D308A | 0.5 | 0.2 |
| D308S | 1§ | 0.2 |
| H309S | 0.1§ | <0.00004* |

**Supplementary Table 2. Relative binding and incision activities of APE1 mutant proteins.** Binding and cleavage of double-stranded DNA containing a THF abasic-site analog were measured as previously described.[34-39] All activities are reported relative to wild-type APE1 (set to 1).

### New measurements corresponding to results presented in Figure 2.

§ measurements not previously reported.

* Indicates limitations of the radiolabeled substrate assay.

n.d., not determined.

#### SUPPLEMENTARY MOVIES

**Supplementary Movie 1. Structural features of APE1 underpinning its proficient $Mg^{2+}$-dependent catalytic mechanism.** The APE1–Mg–DNA ternary complex is shown with DNA colored in pink, APE1 colored by secondary structure (helices in violet, □-sheets in yellow, loop regions in silver). Active site residues are shown explicitly in stick representation and colored by atom type (C in gold, N in blue, H in white, O in red, Mg in green). Zoomed-in views of the active site are shown in stick representation for BO-AIMD derived states S1, S2 and S3, respectively. Active site residues are labeled and coordination of the Mg ion to the oxygen atoms of the scissile phosphate is shown as black dashed lines.

**Supplementary Movie 2. Conformational rearrangements in the APE1 active site during the state S1 to S2 conformational transition.** Active site residues are shown explicitly in stick representation and colored by atom type (C in gold, N in blue, H in white, O in red, Mg in green). Active site residues directly involved in mechanism are labeled. Distances from the scissile phosphate to the nucleophilic water molecule and the Mg ion are shown as black dashed lines.

**Supplementary Movie 3. Conformational rearrangements in the APE1 active site during the state S2 to S3 reactive transition.** Active site residues are shown explicitly in stick representation and colored by atom type (C in gold, N in blue, H in white, O in red, Mg in green). Active site residues directly involved in mechanism are labeled. Distances between the scissile phosphate, the nucleophilic water molecule, and the Mg ion are shown as black dashed lines.
